# Synergistic interactions of quercetin with antibiotics against biofilm associated clinical isolates of *Pseudomonas aeruginosa in vitro*

**DOI:** 10.1101/601336

**Authors:** C. Vipin, M. Mujeeburahiman, K. Saptami, A.B. Arun, P.D. Rekha

## Abstract

Development of extreme resistance to multiple antibiotics is the major concern in infections due to biofilm forming *Pseudomonas aeruginosa.* The existing antibiotics have become ineffective against biofilm associated infections and hence, in this study, the combinatorial efficacy of antibiotics with a quorum sensing inhibitor (quercetin) was tested against biofilm forming *P. aeruginosa* isolates. The effect of drug combinations was studied by the checkerboard method. The fractional inhibitory concentration index (FICI) was calculated for determining the synergistic effect. Additionally, biofilm cell viability, time-kill and live-dead assays were performed to study the combinatorial effect. MIC of quercetin against all the *P. aeruginosa* strains was 500 μg/mL. However, quercetin at 125 μg/mL showed synergistic effect with ½ × MIC or ¼ × MIC of all the antibiotics against all the strains. Quercetin (125 μg/mL) with ½ MIC of levofloxacin and tobramycin combinations were highly effective with ≥80% killing of biofilm associated cells. Increasing the concentration to 250 μg/mL with ½ × MIC antibiotics could completely inhibit the biofilm cell viability in quercetin combination with amikacin and tobramycin. The findings show that quercetin combinations can enhance the treatment outcome against *P. aeruginosa* infection and this approach may reduce antibiotic overuse and selection pressure.

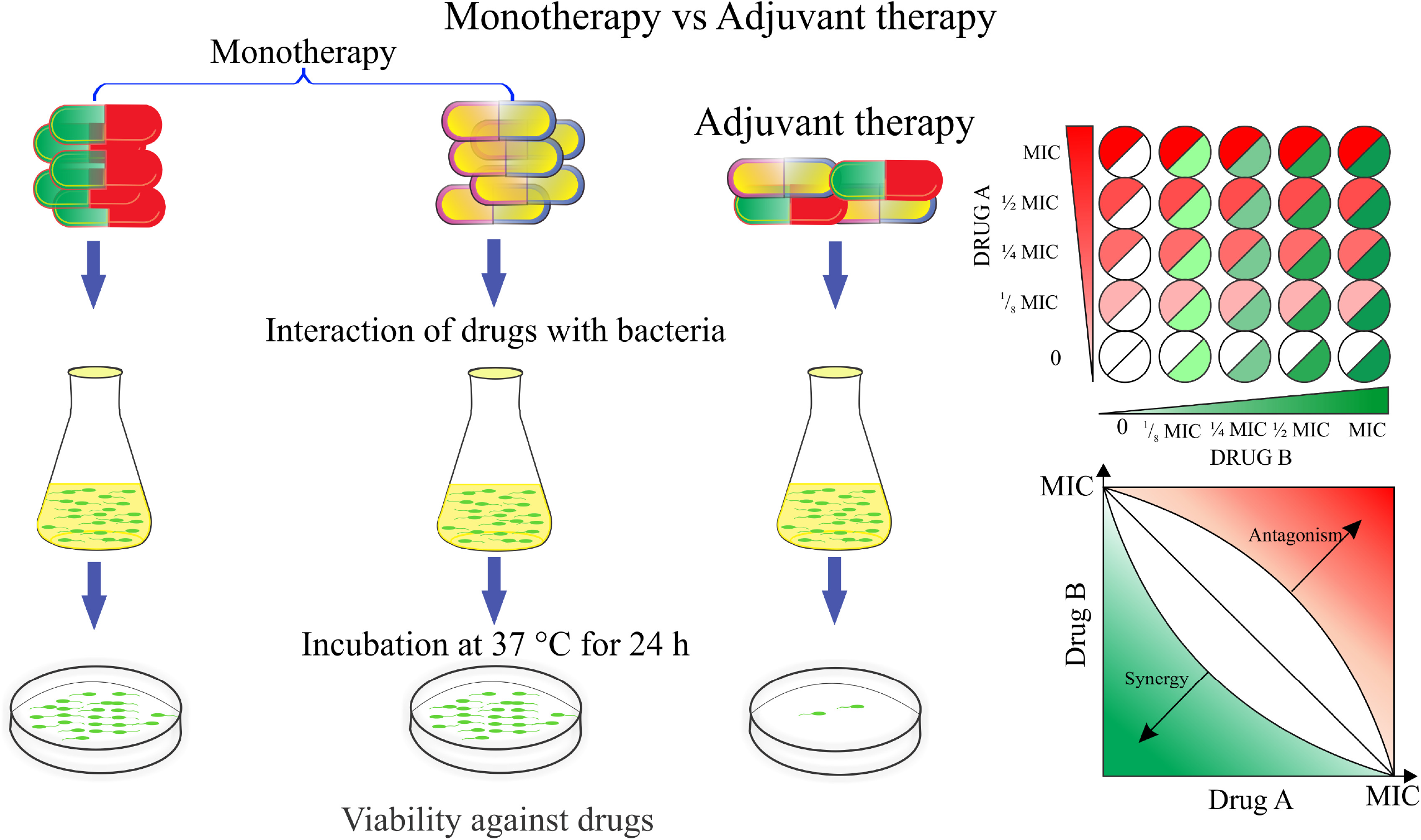
Graphical abstract.

## Introduction

The emergence of resistance to the antibiotics among the biofilm forming *Pseudomonas aeruginosa* has become a great public health problem worldwide. Infection by *P. aeruginosa* is complicated due to its various virulence factors, such as extracellular proteases, toxins and the formation of biofilm (Neidig *et al.*, 2013; Newman *et al.*, 2017). The emergence of antibiotic resistance is believed to be due to the overuse of drugs that allows the bacteria to adapt and modify the virulence factors (Winstanley *et al.*, 2016). The important mechanisms to resist different classes of antibiotics are; mutations in target genes such as DNA gyrase and topoisomerase IV; over-expression of efflux pumps; changes in the cell envelope; downregulation of membrane porins (Kamath *et al.*, 2016), modified lipopolysaccharide component of the outer cell membrane; production of beta-lactamase and aminoglycoside modifying enzymes. Biofilm formation is another trait that offers benefit to the bacteria against antibiotic stress and hence, the cells inside the biofilms are more resistant to antibiotics than the planktonic cells (Melander, 2010; Olsen, 2015). Because of this versatility of bacteria, the existing antibiotic treatments are often ineffective against *P. aeruginosa* persistent infections. Colistin, a polymyxin antibiotic is considered as the last resort therapy for multi-drug resistant (MDR) *P. aeruginosa*, but due to its extreme cytotoxicity, its use is restricted. However, the emergence of colistin resistant strains are reported from various parts of the world warranting serious measures (Nation and Li, 2009; Srinivas and Rivard, 2017). Hence, efforts towards minimizing the colistin usage require alternative strategies and more research into developing effective combinatorial approaches.

A cell to cell communication system called quorum sensing regulates the expression of virulence factors, biofilm formation and antibiotic resistance in *P. aeruginosa.* It is believed that the inhibitors of quorum sensing enhance the drug sensitivity of the existing antibiotics (Brackman *et al.*, 2016). Quercetin, a dietary flavonoid has quorum sensing inhibitory activity (Vasavi *et al.*, 2016) and can inhibit the virulence factors and biofilm formation in *P. aeruginosa* (Paczkowski *et al.*, 2017) with low bactericidal activity (Sakharkar *et al.*, 2009). It is also known for its antioxidant, anti-obesity, anti-carcinogenic and anti-viral potential. Previous studies have also shown that the quercetin is having cytoprotective activity against *P. aeruginosa* infection (Vipin *et al.*, 2019). Owing to its beneficial and anti quorum sensing properties, in this study quercetin was tested in combination with antibiotics to increase the efficacy of treatment against *P. aeruginosa* isolates *in vitro.*

## Materials and methods

### Bacterial strains, culture conditions and minimal inhibitory concentration

The clinical isolates, *P. aeruginosa* YU-V10 and YU-V28 previously isolated from the urinary catheter of the patients and identified by 16S rRNA gene sequencing were used in the study. *P. aeruginosa* PAO1 was used as reference strain in all the experiments. The isolates were maintained in Tryptic Soya Broth (TSB) at 37 °C. For testing of the minimum inhibitory concentrations (MIC) of antibiotics, Muller Hinton broth (MHB) was used. The antibiotics, microtiter plate (96 well-TPG96), TSB and MHB were purchased from HiMedia (Mumbai, India). Quercetin was purchased from Sigma Aldrich (USA). The MIC of antibiotics (amikacin, levofloxacin, tobramycin, gentamycin and ceftriaxone) and quercetin against *P. aeruginosa* isolates were determined by broth microdilution method using polystyrene-96 well plates according to CLSI guidelines (CLSI, 2014). Briefly, fresh MHB containing antibacterial agents at increasing concentrations, amikacin (0.5 to 64 μg/mL), levofloxacin (0.5 to 32 μg/mL), tobramycin (0.5 to 32 μg/mL), gentamycin (0.5 to 32 μg/mL), ceftriaxone (0.5 to 32 μg/mL) and quercetin (62.5 to 1000 μg/mL) were inoculated with overnight cultures of the bacteria at a cell density of 1×10^5^ CFU/mL and incubated at 37 °C for 24 h. After incubation, the plates were observed for the growth. MIC was interpreted as the lowest concentration of the antibiotic at which no visible growth was observed.

### Checkerboard assay and fractional inhibitory concentration

The interaction of two antimicrobial agents was investigated by the checkerboard method using 96-well microtiter plates. Amikacin, levofloxacin, ceftriaxone, gentamycin and tobramycin were tested in combination with quercetin respectively against all the isolates of *P. aeruginosa* at different fractional MIC concentrations (½ × MIC, ¼ × MIC and ⅛ × MIC). The concentrations of quercetin tested were 62.5 μg/mL (⅛× MIC), 125 μg/mL (¼ × MIC) and 250 μg/mL (½ × MIC). Antibiotic stocks of 10 mg/mL concentration were prepared and diluted to the required concentrations using fresh MHB. Quercetin stock (10 mg/mL) was prepared in dimethyl sulfoxide (DMSO). Bacterial inoculum was prepared from the overnight cultures of the isolates by centrifugation and resuspending in fresh MHB. The antibiotic solution was added to the wells to attain the concentrations of individual antibiotic in combination and inoculated with the bacteria (100 μL) to an initial density of 1×10^5^ CFU/mL. The contents were incubated at 37 °C for 24 h and observed for visible turbidity. The antimicrobial agents alone at MIC and sub MIC concentrations (½ × MIC, ¼ × MIC and ⅛ × MIC) were kept for comparison. The MIC of the drug combinations was defined as the concentration of no visible growth after 24 h incubation at 37 °C. The fractional inhibitory concentrations (FICs) and FIC index (FICI) were calculated as follows (Louie *et al.*, 2011);

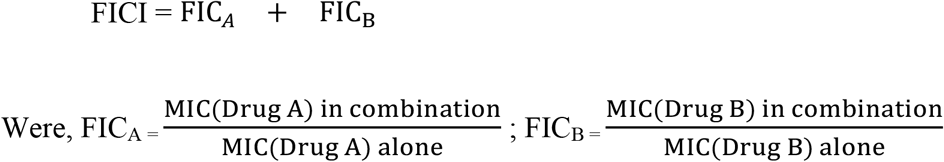

Synergy was defined based on the FICI and interactions were classified as follows: FICI ≤0.5 as synergistic; >0.5 to ≤1 as an additive; and FIC index >1 as antagonistic (Mulyaningsih *et al.*, 2010).

### Time-kill assay

Time-kill assay was performed to study the combined effects of antimicrobial agents on the bacterial growth at different time points (0-12 h). For this, the 96 well plates containing drug combinations and inoculum were prepared as described in the previous section. The plates were incubated at 37 °C and the bactericidal effect of antibiotic and quercetin combinations were evaluated at every 0 to 12 h intervals. The growth was measured by recording OD_600_ spectrophotometrically (FLUOstar Omega, BMG Labtech, Germany) (Pournaras *et al.*, 2011).

### Evaluation of drug combinations on biofilm cell viability

For the development of biofilm, the test strains were incubated in 96-well plates containing TSB media with an initial cell density of 1 × 10^5^ CFU/mL for 24h at 37 °C statically. Following the incubation, the planktonic cells were removed by decanting the culture broth. The adherent biofilm was washed gently with sterile PBS and different combination of antibiotics with quercetin was added. The volume in each well was adjusted to 200 μL using fresh TSB. The plates were incubated for 8 h at 37 °C under the static condition to see the effect of drug combinations on the biofilm associated cells. After incubation, the liquid contents were discarded and washed with sterile PBS in order to remove the non-adherent cells. To see whether there were residual cells in the biofilm, the biofilm viability was tested by adding fresh TSB to allow the growth of residual cells for 24 h at 37 °C. After incubation, the growth was measured at OD_600_ and compared with the growth in the non-treated control wells to calculate the percent cell viability in the biofilm. Further, the samples from each well were serially diluted and plated on tryptic soy agar plates to confirm the cell viability (Nair *et al.*, 2016).

### Live/dead assay for the biofilm

The live/dead staining was used to study the biofilm viability after treating with the quercetin (125 μg/mL) with respective ½ × MIC of antibiotics. For these biofilms were developed on glass coupons by incubating in TSB for 24 h at 37 °C using 24 well plates under static condition. The glass coupons containing biofilms were gently removed, washed with sterile PBS and transferred to fresh 24 well plates containing antibiotics in fresh TSB. After 8 h of incubation, the coupons were removed, washed with PBS and stained with a mixture of propidium iodide and acridine orange (1:1) for 15 min at room temperature under dark. The coupons were observed using confocal laser scanning microscope (CLSM, LSM 710, Carl Zeiss, Jena, Germany). The fluorescence emission was measured at 555 nm for acridine orange and 625 nm for propidium iodide. Both dyes were excited at 485 nm. The percentage of live cells in the biofilm was plotted as a function of the ratio between the green and red fluorescence intensities. Images of the biofilm acquired were processed using Zen 2011 software (Carl Zeiss) (Bédard *et al.*, 2014).

### Statistical analysis

The experiments are performed thrice in triplicates. The statistical analysis was performed by student’s test using SPSS version 22 (SPSS Inc., Chicago, IL, USA). P values of <0.05 were regarded as significant.

## Results

### Minimum inhibitory concentrations

The MIC of the antibiotics against YU-V10, YU-V28 and PAO1 is given in **Table. 1**. The MIC for levofloxacin ranged between 2 and 8 μg/mL, tobramycin it was between 3 and 6 μg/mL.

**Table 1:** MIC of antimicrobials tested against clinical isolates of *P. aeruginosa*

| Strains | MIC (µg/mL) |  |  |  |  |  |
| --- | --- | --- | --- | --- | --- | --- |
|  | Levofloxacin | Ceftriaxone | Gentamycin | Tobramycin | Amikacin | Quercetin |
| PAO1 | 2 | 8 | 4 | 3 | 8 | 500 |
| YU-V10 | 4 | 8 | 4 | 3 | 8 | 500 |
| YU-V28 | 8 | 8 | 4 | 6 | 8 | 500 |

### Interaction of quercetin with antibiotics

In combination with quercetin, the antibiotics showed 2 to 4-fold decrease in the MICs. The quercetin – antibiotic combinations showed synergistic effect against all the tested isolates with FICI ranging from 0.25 to 0.5. In combination therapy, quercetin at a concentration of 62.5 μg/mL (⅛ × MIC) could decrease the antibiotics dose to a minimum of ½ × MIC. Quercetin at the same concentration (62.5 μg/mL) could reduce the MIC of amikacin by ⅛ and ceftriaxone by ½ with FICI ranging from 0.25 to 0.375 against different isolates. Quercetin at a concentration of 125 μg/mL (¼ × MIC) showed synergism with gentamycin (⅛ × MIC) and levofloxacin (¼ and ⅛ × MIC) with FICI of 0.375 to 0.5 against all the tested isolates. The checkerboard test results and isobolograms are given in figure **Fig. 1a-d**. The synergy between quercetin and all the tested antibiotics against *P. aeruginosa* isolates are given in **Table. 2**.

**Fig 1.**
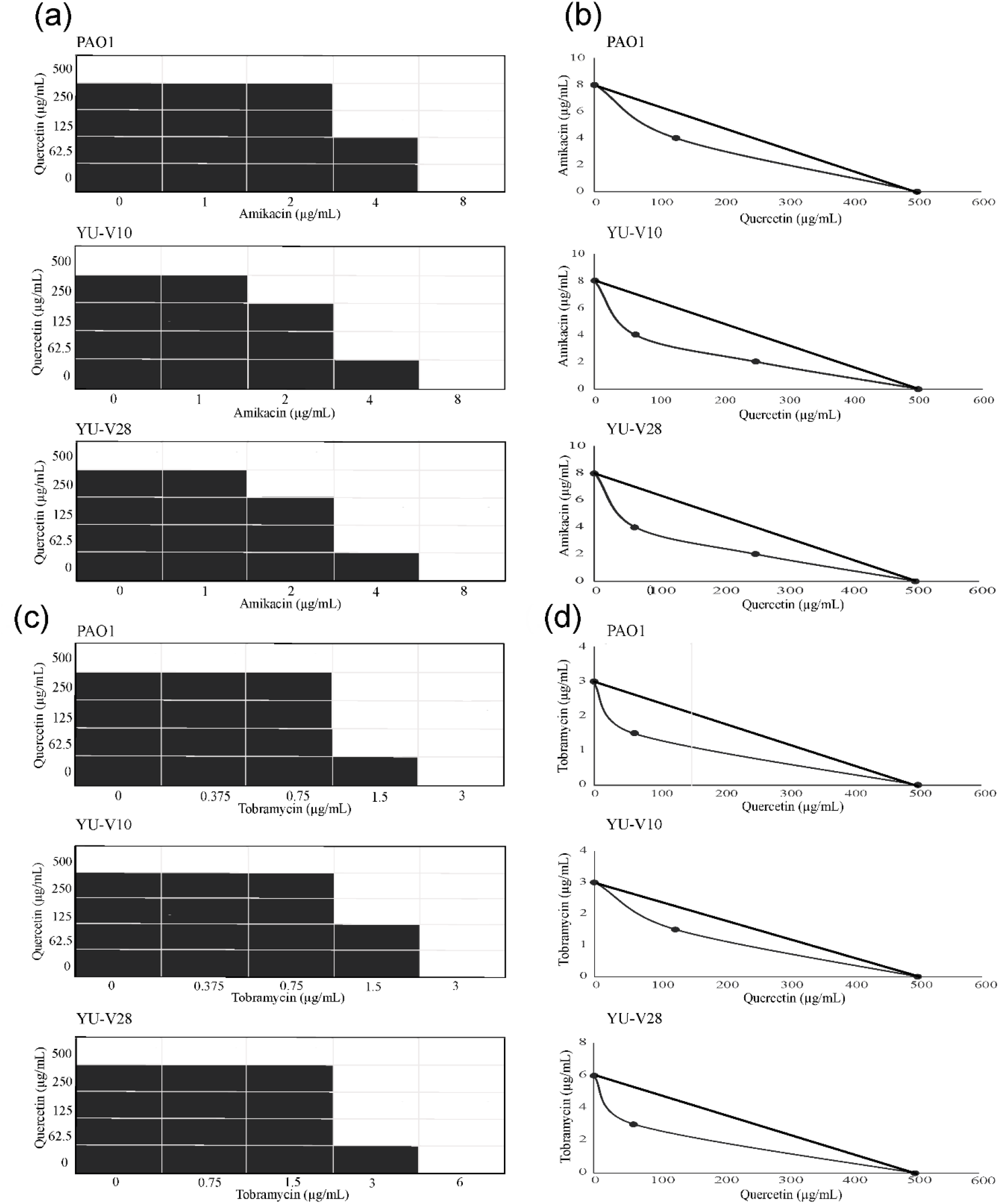
Synergistic effect between quercetin with antibiotics against *P. aeruginosa* isolates as determined by the checkerboard assay and calculation of the fractional inhibitory concentration (FIC) index. Checkerboard assay results and isobolograms: (a,b) quercetin/amikacin (FIC index = 0.25); (c,d) quercetin/ tobramycin (FIC index = 0.25 for YU-V10 and PAO1; 0.375 for YU-V28). shading: visible growth

**Table 2:**
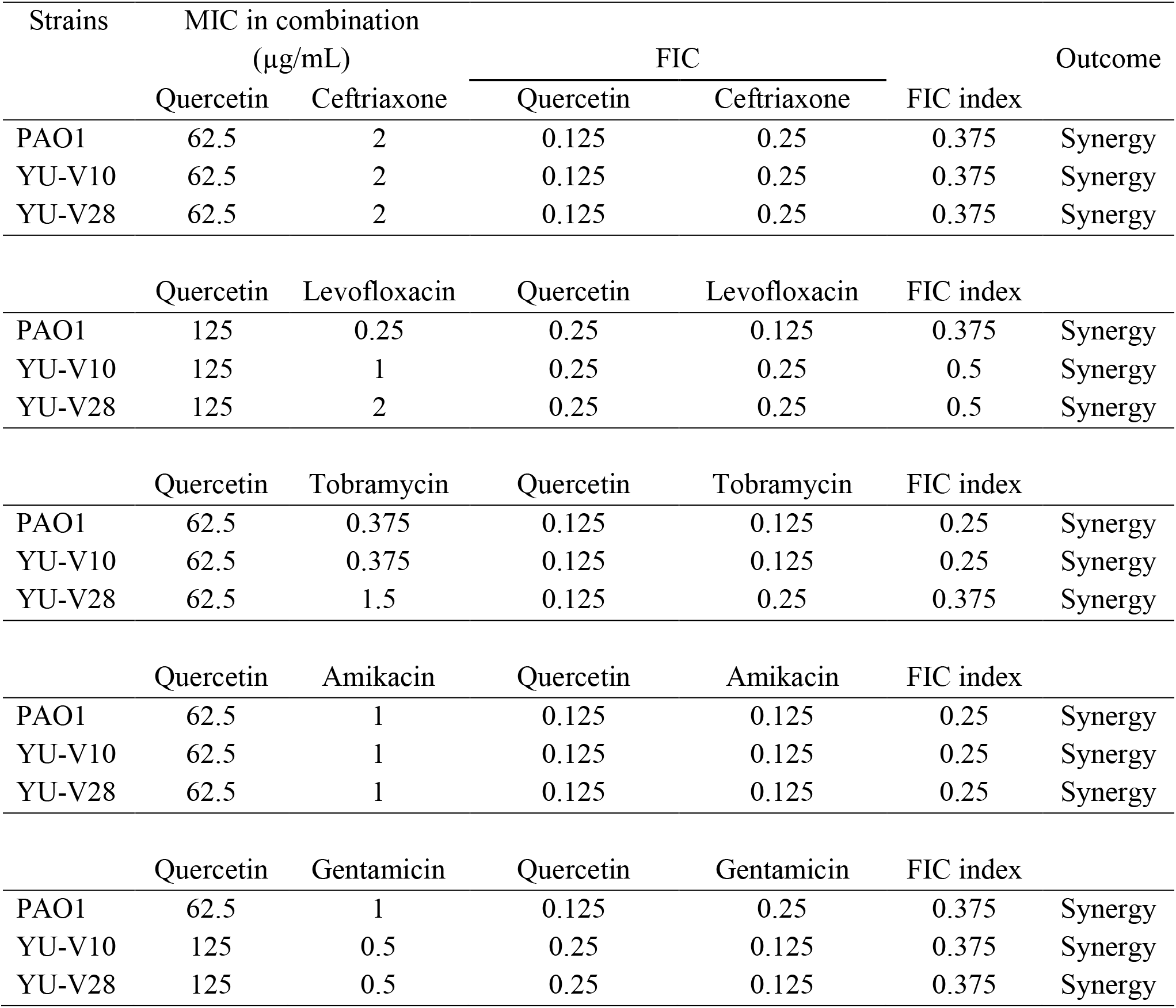
The synergistic effect of antibiotics in combination with quercetin (FICI)

### Effect of drug combinations on growth and biofilm viability

In the time kill assay, the effect of drug combinations on bacteria in the early stage of growth showed a significant reduction in the bacterial growth compared to control. The combination of quercetin (62.5 μg/mL) with ceftriaxone (¼ × MIC), tobramycin (¼ × MIC), amikacin (⅛ × MIC) showed synergism with the higher killing rate. All the strains showed a significant reduction in growth (p<0.05) in tested combinations at 0-12 h compared to respective untreated controls (**Fig. 2a-b**).

**Fig 2.**
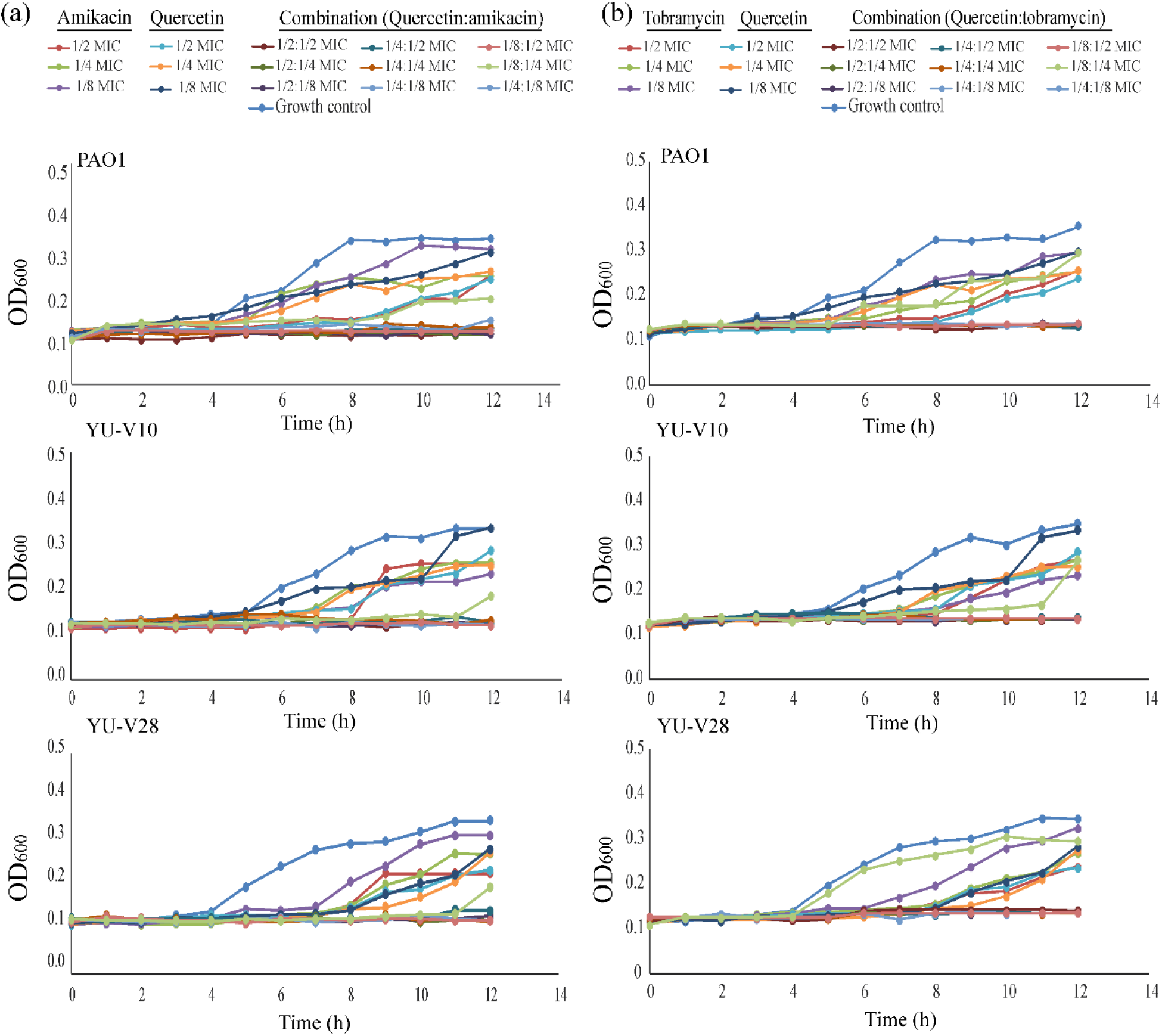
Combinatorial effect of drugs against *P. aeruginosa* isolates in growth initiation at 37 °C for 0-12 h incubation. a) quercetin and amikacin, b) quercetin and tobramycin. The combinatorial treatment significantly reduced growth initiation by *P. aeruginosa*.

The efficacy of drug combinations on the viability of a fully formed biofilm community is given in **fig 3 and 4**. The ½ × MIC dose of antibiotics with quercetin at a concentration of 125 μg/mL (¼ × MIC) showed ≥80% reduction in biofilm cell viability. Quercetin at a concentration of 250 μg/mL (½ × MIC) with ½ × MIC of antibiotics completely inhibited the biofilm cell viability in PAO1 in quercetin-amikacin and quercetin-tobramycin combinations (**Fig. 3a-b**). The live/dead assay also confirmed the decrease in the viable cells in the biofilm matrix with drug treatment. The CLSM observations of biofilm matrix treated with quercetin (125 μg/mL) in combination with ½ × MIC of antibiotics showed synergistic activity on biofilm matrix with fluorescent intensity ratio ranging from 53-100% which corresponds to the percent of dead cells in the biofilm matrix (**Fig. 4**).

**Fig 3.**
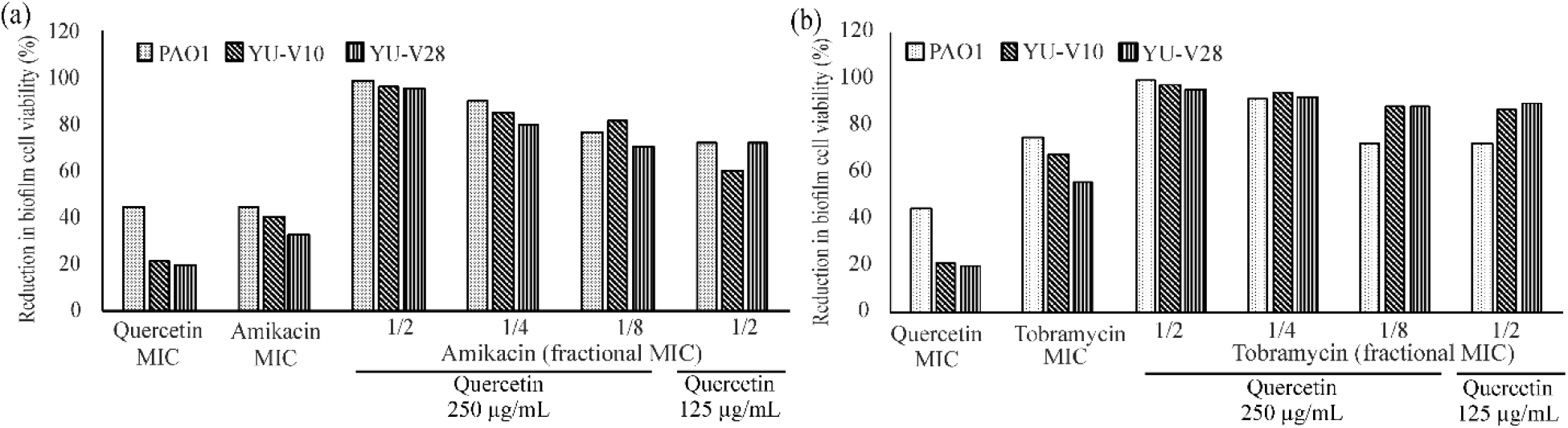
Effective combination of antibiotics with quercetin showing inhibition of the biofilm cell viability. Preformed biofilms were treated with synergistic drug combinations, (a) quercetin-amikacin, (b) quercetin-tobramycin combinations.

**Fig 4.**
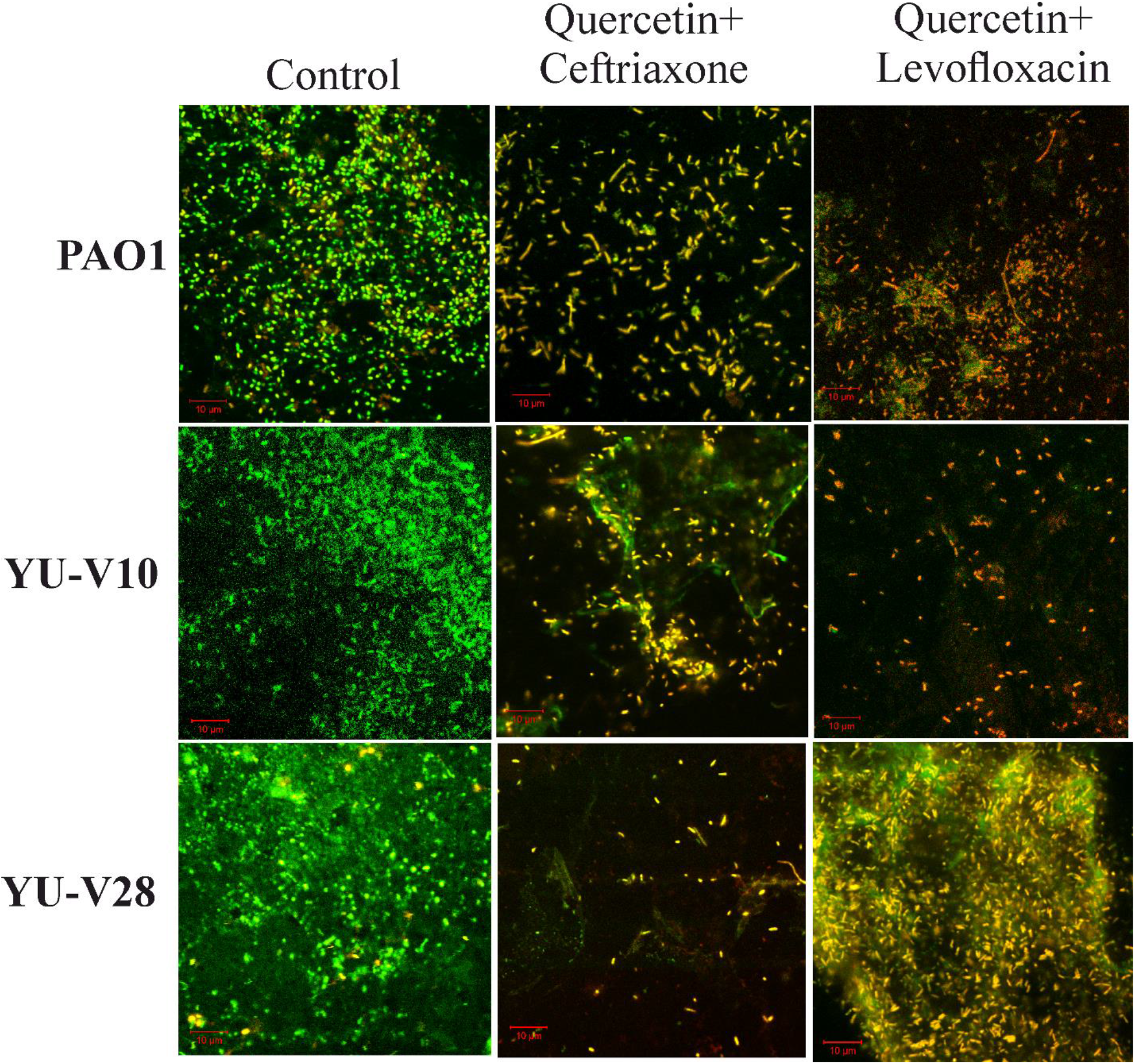
Synergy between quercetin and antibiotics. Representative images of live/dead staining of mature *P. aeruginosa* (YU-V10, YU-V28 and PAO1) biofilm treated with quercetin (125 μg/mL) and ½ ×MIC ceftriaxone, quercetin (125 μg/mL) and ½ ×MIC levofloxacin compared with non-treated control. Green colour indicates the live cells and yellow to red colour indicates dead cells.

## Discussion

This study showed that the quercetin in combination with antibiotics can increase the antibacterial and antibiofilm efficacy against biofilm forming *P. aeruginosa* isolates. Quercetin is a phytocompound with anti-quorum sensing potential against *P. aeruginosa* (Vasavi *et al.*, 2014). In *P. aeruginosa*, quorum sensing plays a significant role in biofilm formation as well as controlling the expression of virulence factors. The anti-quorum sensing compounds disrupt the bacterial community coordination by blocking the quorum sensing system and regulating the population density. This study showed that the most effective quercetin – antibiotic combination was quercetin-tobramycin. Quercetin at a concentration of 62.5 μg/mL with 0.375 μg/mL of tobramycin could inhibit the bacterial growth and biofilm formation at the lowest dose. It has been reported that quercetin can be consumed up to 5 g daily without any adverse effects (Lu *et al.*, 2016). Previous studies have also shown that the quercetin does not have major adverse effect on normal human cells (Vizzotto *et al.*, 2014). Quercetin is a prominent constituent of dietary supplements and quercetin-based supplements are also recommended for the prevention of cancer, improvement of cardiovascular functions and mental health (Jain *et al.*, 2016). The anti-inflammatory properties of quercetin can also benefit patients by providing antiinflammatory effect against the inflammation caused by the infection. The inflammation caused due to infection can lead to fever, erythema, oedema, pain and loss of function (Basil and Levy, 2015). Quercetin along with antibiotics will reduce such inflammatory responses along with targeting the pathogens.

At MIC concentrations of the antibiotics, the biofilms are difficult to be eradicated. It requires 4 ×MIC concentration to effectively remove the biofilm. The quercetin – antibiotic combinations showed synergistic effect on biofilm cell viability which was significantly higher than the monotherapy, which is an advantage as current strategies are often inefficient to kill or restrict the growth of the cells in the biofilm. The synergy of drug combination is due to the antiquorum sensing mediated biofilm inhibition by quercetin. Anti-quorum sensing activity and existing antibiotics can generate synergistic effects and broaden the antimicrobial effectiveness. The combinatorial treatment showed minimum of 80% decrease in live cells in the biofilm matrix by the effective combinations with 8 h of treatment. However, in the respective monotherapy MIC dose, it was 19.4 to 74.4% reduction in viability in different isolates. This indicates that the combined therapy is effective in penetrating to the biofilm matrix and causing cell death or even depletion.

Apart from anti quorum sensing activity, quercetin can damage the bacterial cell wall ultrastructure and cell membrane integrity of the bacteria (Wang *et al.*, 2017). The increase in cell membrane permeability makes the combination treatment more effective. Quercetin can also modulate the bioavailability of drugs and thus increase the efficacy of the antibiotics (Schutte *et al.*, 2008; Umathe *et al.*, 2008). The drugs having bioavailability enhancing properties can make the target cells more susceptible to drugs. The use of phytocompounds combined with existing antibiotics is a promising strategy to combat antibiotic resistance and biofilms associated *P. aeruginosa* infections. The findings of this study provide evidence that quercetin has the potential to target biofilm forming *P. aeruginosa* in combination with antibiotics. Quercetin is non-toxic in nature; hence quercetin can be used safely in combination with suitable antibiotics against *P. aeruginosa* isolates after appropriate *in vivo* studies.

## Ethical approval

The study was approved by the institutional ethics committee of Yenepoya University with protocol number: YUEC386/2016.

### Informed consent

Informed consent was obtained from all individual participants included in the study

